# Calcium dobesilate reduces VEGF signaling by interfering with heparan sulfate binding site and protects from vascular complications in diabetic mice

**DOI:** 10.1101/661793

**Authors:** Florence Njau, Nelli Shushakova, Heiko Schenk, Vera Christine Wulfmeyer, Robin Bollin, Jan Menne, Hermann Haller

**Affiliations:** Division of Nephrology, Hannover Medical School, Hannover, Germany

**Keywords:** Calcium dobesilate, VEGF, VEGFR-2, Heparan sulfate, diabetic nephropathy

## Abstract

Inhibiting vascular endothelial growth factor (VEGF) is a therapeutic option in diabetic microangiopathy. However, VEGF is needed at physiological concentrations to maintain glomerular integrity; complete VEGF blockade has deleterious effects on glomerular structure and function. Anti-VEGF therapy in diabetes raises the challenge of reducing VEGF-induced pathology without accelerating endothelial cell injury. Heparan sulfate (HS) can act as a co-receptor for VEGF. Calcium dobesilate (CaD) is a small molecule with vasoprotective properties that has been used for the treatment of diabetic microangiopathy. Preliminary evidence suggests that CaD interferes with HS binding sites of fibroblast growth factor. We therefore tested the hypotheses that (1) CaD inhibits VEGF signaling in endothelial cells, (2) that this effect is mediated via interference between CaD and HS, and (3) that CaD ameliorates diabetic nephropathy in a streptozotocin-induced diabetic mouse model by VEGF inhibition. We found that CaD significantly inhibited VEGF_165_-induced endothelial cell migration, proliferation, and permeability. CaD significantly inhibited VEGF_165_-induced phosphorylation of VEGFR-2 and suppressed the activity of VEGFR-2 mediated signaling cascades. The effects of CaD in vitro were abrogated by heparin, suggesting the involvement of heparin-like domain in the interaction with CaD. In addition, VEGF_121_, an isoform which does not bind to heparin, was not inhibited by CaD. By applying proximity ligation assays to endothelial cells, we show inhibition of interaction in situ between HS and VEGF and between VEGF and VEGFR-2. Moreover, CaD reduced VEGF signaling in diabetic kidneys and ameliorated diabetic nephropathy and neuropathy, suggesting CaD as a VEGF inhibitor without the negative effects of complete VEGF blockade and therefore could be useful as a strategy in treating diabetic nephropathy.

## Introduction

Diabetic nephropathy is one of the most important microvascular complications of diabetes mellitus and is responsible for 40-50% of all cases of end-stage renal disease (ESRD), despite various treatment strategies, such as intensive blood glucose control (1,2), lowering of blood pressure (3,4) or renin-angiotensin-system blockade (5) that have been established over the last 20 years (6,7). The complex pathogenesis of diabetic nephropathy makes the development of evidence-based therapeutic strategies difficult (8).

An increased expression of vascular endothelial growth factor (VEGF) has been observed in animal models and in diabetic patients (9–11). Increased VEGF-A/VEGFR-2 signaling contributes to renal disease in several important ways, including vascular permeability (12), vasodilatation, hyperfiltration (13,14), capillary growth, and monocyte chemotaxis (15,16). Inhibiting VEGF seems to prevent the development of nephropathy in animal models. Treatment with an anti-VEGF_165_ antibody results in a significant attenuation of albuminuria in diabetes animal models (13). This effect was confirmed in other diabetes models (1,17). However, anti-VEGF treatment in the prevention of microvascular disease is associated with serious obstacles, since, for example, VEGF_165_ antibodies cause renal damage and hypertension in patients, and proteinuria commonly occurs after anti-VEGF therapy (18). VEGF has been observed to have an important role in maintaining the endothelial integrity, given that anti-VEGF in patients as well as conditional ablation of VEGF in adult mice led to microangiopathy (19,20). These conflicting observations have led to the hypotheses that on-the-one hand under physiological conditions VEGF signaling is necessary to maintain endothelial stability, while on-the-other hand, overexpressing VEGF and its signaling, as occurs in diabetes, leads to endothelial damage and microvascular disease.

Calcium dobesilate (CaD) is a small molecule which has been used in Asia and South America to treat various vascular disorders including diabetic microvascular disease, for years. CaD also seems effective for the treatment of rosacea and psoriasis (21,22), which are also associated with increased VEGF activity. However, the pharmacology of CaD is poorly understood. CaD belongs to the 2,5-dihydroxyphenylic acids, a newly described family of molecules which interfere with growth factor signaling (23), CaD binds to the heparin-binding domain of FGF-1, thus reducing FGF-1 activity (23). We reasoned that CaD could function as a novel VEGF antagonist. We used cultured endothelial cells and animal models and found that CaD indeed reduces exaggerated VEGF signaling, while maintaining physiological effects of VEGF. The 2,5-dihydroxyphenylic-acid compound class could represent a novel VEGF antagonist without adverse side effects.

## Materials and Methods

### Materials

Primary human umbilical vein endothelial cells (HUVECs; ATCC®PCS-100.010) were purchased from ATCC (Wesel, Germany) and cultured in EGM™BulletKit™without exogenous VEGF (Lonza). CCK-8 cell viability assay kit (Dojindo Molecular Technologies, Munich Germany), polycarbonate filters (ThinCert^™^, Greiner bio-one). All VEGF-A used in this study were VEGF_165_ isoform unless designated otherwise. The recombinant VEGF_165_, VEGF_121_ and biotinylated-VEGF_165_ (bt-VEGF_165_), VEGFR-1, VEGFR-2 and recombinant human Heparanase were from R&D Systems Inc. (Wiesbaden-Nordenstadt, Germany). Heparin sodium salt from porcine intestinal mucosa and Calcium dobesilate (2,5-Dihydroxybenzenesulfonic acid calcium salt), Fluorescein isothiocyanate-dextran, molecular mass: 70 kDa, Duolink® In Situ PLA kit and probes, calcein-AM and streptozotozin (STZ) were from Sigma Aldrich (Taufkirchen, Germany). Rabbit primary antibodies for VEGFR-2, pTyr1175, p-ERK1/2, ERK, pP38, pMEK and MEK were acquired from Cell Signaling Technology (Leiden, The Netherlands) and F4/80 (clone A3-1; BioLegend, San Diego, CA, USA). Mouse anti-heparan sulfate proteoglycan (mAb F58-10E4) was from Amsbio. GAPDH and all secondary antibodies (except Cy3, Jackson ImmunoResearch, West Grove, USA) were from Santa Cruz Biotechnology (Heidelberg, Germany). Streptavidin-HRP (Thermo Fisher Scientific) and phalloidin-Alexa fluor 488 was from Invitrogen, Carlsbad, CA, USA. Other primary antibodies were obtained as indicated; Occludin (Invitrogen), Claudin-5 (Bioworld Technology, Inc), ZO-1 (BD Transduction Laboratories), Vinculin (Chemicon). VEGF quantikine ELISA kits and goat anti-human VEGF_165_ antibody (AF-293-NA) were from R&D Systems. Phospho-VEGFR-2 (Tyr1175) Sandwich ELISA Kit was from Cell Signaling Technology. Mouse albumin ELISA kit (Bethyl Lab, Hamburg, Germany), paraformaldehyde (Merck, Darmstadt, Germany), histoclear (Biozym, Hessisch Oldendorf, Germany). VectaShield mounting medium (Vector Laboratories Inc., Burlingame, CA). Periodic acid (0.5%) and Schiff’s reagent (Merck), hematoxylin was from Fluka.

### Animal model

Male 10 weeks old 129/SV mice (Charles River, Germany) were held in individually ventilated cages and received a standard diet with free access to tap water. Weight-matched 129/SV mice received either 125 mg/kg body weight STZ (Sigma-Aldrich) in 50 mM sodium citrate (pH 4.5) or sodium citrate buffer (nondiabetic control group) intraperitoneally on day 1 and 4 for induction of hyperglycemia (glucose > 15 mmol/l). Mice received no insulin during the study. All procedures were performed according to the guidelines from the Federation of European Laboratory Animal Science Associations and were approved by local authorities (Lower Saxony State Departments for Food Safety and Animal Welfare); approval number 33.19-42502-04-15/1925.

Diabetic mice (n=20) were treated with placebo (saline), 100 mg/kg body weight CaD, or 30 mg/kg body weight enalapril by gavage. Body weight and glucose levels were measured every week. HbA1c (Olympus AU400) and Kidney function (serum creatinine) were measured at 6 and 12 weeks. Albuminuria was assessed using a mouse albumin ELISA kit (Bethyl Lab, Hamburg, Germany).

Sensory nerve conduction velocity (NCV) studies were performed in mice anesthetized with 2% isoflurane at week 6 and 12. Tail sensory NCV was determined by stimulating proximally along the tail at a recorded distance of 3 cm. For the measurement, a neuro-screen from Toennies Inc. was used. After 12 weeks of follow up the mice were euthanized by expose to isoflurane at 5% concentration which was continued for 1 min after breathing stop. Thereafter bilateral thoracotomy and laparotomy were performed and kidneys were perfused with ice cold saline solution via the left heart ventricle.

### Histology and immunohistochemistry

Histological and morphometric analyses were carried out on 3% PFA-fixed paraffin sections (2 µm), stained with periodic acid-Schiff (PAS) reaction and alcian blue.

For immunofluorescence, paraformaldehyde-fixed and paraffin-embedded tissue sections (2μm) were processed as previously described (24). After blocking with 10% rabbit serum, paraffin sections were stained with antibodies against pP38 and F4/80 and with a secondary antibody conjugated to Cy3. Specimens were analyzed using a Zeiss Axioplan-2 imaging microscope with AxioVision 4.8 software (Zeiss, Jena, Germany).

### Cell culture

Primary human umbilical vein endothelial cells (HUVECs) were routinely cultured in 0.1% gelatin pre-coated flasks or dishes, up to passage 6. The effect of CaD alone (0, 10, 20, 50, 100, and 200 µM) on cellular viability was assessed by CCK-8 kit using a Tecan Microplate Reader (Genios). To measure the effect of CaD on VEGF-induced cell viability, HUVECs (1 × 10^4^ cells/well) were treated with VEGF (20ng/ml) pre-mixed with various concentrations of CaD (0, 50, 100 and 200 µM) in starvation medium for 24 h and 48 h. The number of viable cells is presented relative to untreated controls.

### Wound healing

Confluent HUVECs monolayer was scraped using a 0.2ml pipette tip after 2 h of complete serum starvation. Subsequently, cells were washed; fresh EGM medium containing 0.5% FCS and different concentrations of CaD (0, 50 and 100 µM) with or without 20 ng/mL VEGF was added. Images were taken using a Leica DM 14000B microscope after 16 h incubation. The gap distance of migrated cells was quantitatively evaluated using ImageJ software.

### Endothelial cell transwell invasion assay

The motility of HUVECs was performed in 24-well transwell plates using 8μm polycarbonate filters coated with 0.1% gelatin. Cells were seeded into the upper chambers at a density of 1 × 10^5^ cells per chamber, the bottom chambers were filled with 600μL 0.5% FCS EGM supplemented with VEGF (20 ng/mL) with or without CaD (0, 50, 100 µM). After 24 h, the number of migrated cells was evaluated with calcein-AM using Greiner bio-one quantitative cell migration assay protocol. The results were the means from 3 replicates of each experiment.

### Endothelial permeability assay

Permeability across endothelial cell monolayers was measured using gelatin-coated Transwell ThinCert™ 0.4 μm pore size polycarbonate filter in 24 well as previously described (25).

### Western blotting

Cells were seeded into 6 cm dishes till 80-100% confluency and then starved in serum free medium for 2 h. For cells stimulated in the presence of heparin, heparin, CaD and VEGF were premixed and incubated for 1h at 37°C before addition to the cells. For heparinase treatment experiments, cells were starved for 1.5 h then treated for 30 min with heparinase at 37°C, washed 3 times with warm medium before addition of VEGF/CaD mixtures. For the interaction between CaD and VEGFR-2, the cells were first pre-incubated with CaD for 1 h during the starvation time, followed by washing with warm medium. Cells were then exposed to serum free medium supplemented with 25 ng/ml VEGF with or without the indicated CaD concentrations for 2 or 15 min. Cells and mouse kidney tissue were lysed in RIPA buffer and processed as previously described (26).

### F-actin staining and enzyme-linked immunosorbent assay (ELISA)

HUVECs were seeded at 1 × 10^4^ cells/well onto coverslips in a 12-well plate until 60% confluent. Serum-starved cells were treated with or without 100 μM CaD/20 ng/mL VEGF for 15 min. The cells were then fixed in 4% paraformaldehyde and then permeabilized with 0.5% Triton X-100. Actin filaments were stained by phalloidin-Alexa fluor 488 for 1 h at room temperature and nuclei were detected by DAPI. The slides were examined with a Leica DM 14000B confocal microscope. The concentration of VEGF and pVEGFR-2-Tyr1175 in mouse kidney lysates was measured using commercially available ELISA kits.

### Quantitative RT-PCR analysis

mRNA from kidney sections in RNA later was isolated using RNeasy miniprep kit (Qiagen). qPCR was performed on a LightCycler 96 Real-Time PCR System using SYBR Green RT-PCR with the following Quantitec primers from Qiagen; IL-6 (QT0009887), CXCL1 (QT00113253), MCP-1 (QT00167832), IL-1ß (QT01048355) and TNF-α (QT00104006). Quantification was carried out by LightCycler 96 software and the amount of RNA was expressed as fold change relative to the housekeeping gene (β-Actin; QT00095242).

### Solid-phase binding assay of biotinylated VEGF to recombinant human VEGFR1–2

The method was performed as described previously (27). Briefly, 96-well microplate was coated with 500 ng/ml of either VEGFR-1 or −2 in PBS, sealed and incubated overnight at 4°C. After 3 times washes with PBS-Tween 20 (0.05% v/v), the plate was blocked with PBS with 1% (w/v) BSA, and incubated for 2 h at room temperature. After washing, a mixture of bt-VEGF (50 ng/ml) and heparin (1 µg/ml) or various concentrations of CaD in PBS were applied overnight at 4°C. After washing, streptavidin-HRP (1:4000) was added for 2 h, washed and substrate solution was added for 30-45 minutes. Stop solution (2N; H_2_SO4) was added and fluorescence was measured at 450nm.

### Proximity Ligation Assay

HUVECs were treated with VEGF/CaD for 2 min followed by fixation in paraformaldehyde and permeabilization as described above. Blocking and Proximity ligation was performed using a Duolink PLA kit according to the manufacturer’s protocol. To study VEGF/VEGFR2, VEGF/HS and VEGFR2/HS interactions, cells were incubated overnight at 4 °C with goat monoclonal VEGF (1:100) and/or mouse monoclonal heparan sulfate (1:200), rabbit polyclonal anti-VEGFR2 (1:800). All images were taken with a Leica DMI3000 B microscopy with a 20x objective and analyzed with NIH ImageJ software.

### Statistics

All data are expressed as the mean ± SD of indicated *n* values (for *in vitro* data) and mean ± SEM (for *in vivo* data). One-way analysis of variance was used to compare between groups. Data were analyzed using post hoc Bonferroni correction for multiple comparisons. P-values are *(P<0.05), ** (P<0.01), *** (P<0.001).

## Results

### CaD inhibits VEGF-induced VEGFR-2 phosphorylation in HUVECs

After cells were treated with a range of CaD (0-200 µM) for 24 h and 48 h, the cells exhibited no signs of cytotoxicity (data not shown). We then evaluated the inhibitory effect of CaD on VEGF-induced activation of the VEGFR-2 signaling pathway and the angiogenic response in HUVECs. Firstly, CaD was pre-mixed with VEGF_165_ to determine whether CaD binds directly to VEGF molecules. CaD at different concentrations (6-100 µM) was incubated with VEGF (25 ng/mL) for 60 min prior to being added to HUVECs. CaD significantly decreased VEGF-induced VEGFR-2 phosphorylation in a concentration-dependent manner (Fig. 1A). CaD reduced VEGF-induced phosphorylation of VEGFR-2 up to 50% without affecting the overall VEGFR-2 expression level (Fig. 1B, lanes 5 and 6).

**Fig. 1.**
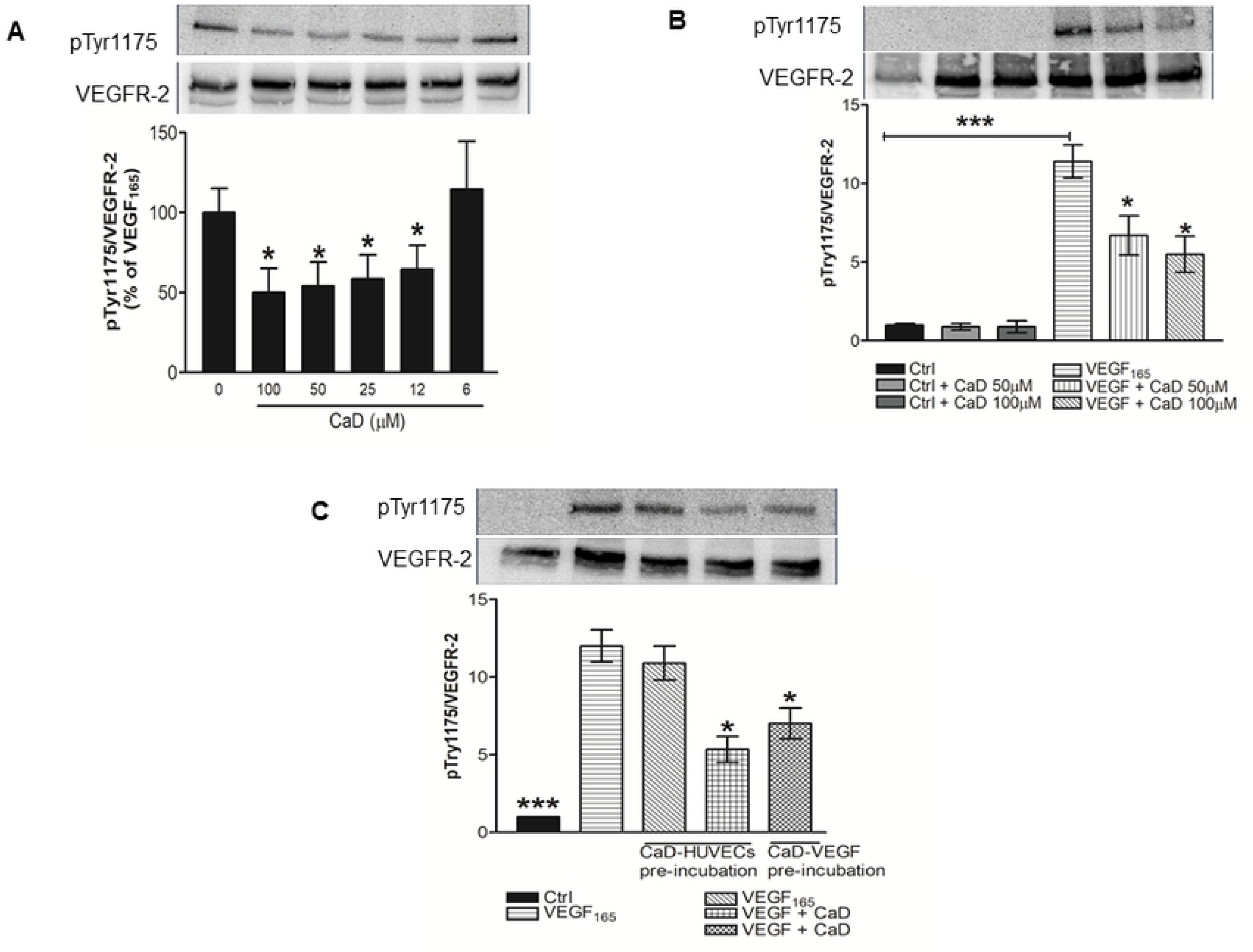
Calcium dobesilate (CaD) inhibits VEGF-induced VEGFR-2 activation. VEGF_165_ (25 ng/mL) was premixed and incubated with: (**A**) various CaD concentrations (6–100 µM) or (**B**) 50-200 µM CaD for 1 h, before exposure to HUVECs for 2 min. (**C**) HUVECs were incubated with 50 and 100µM CaD for 60 min with three subsequent washing steps with warm medium before stimulation with VEGF_165_ for 2 min. Western blot analysis was performed using anti-phospho-VEGFR-2 antibody and total VEGFR2 was used as a loading control after membrane stripping. Each bar represents the mean ± SD (n=3). *P<0.05, ***P<0.001 versus VEGF-treated HUVECs.

Secondly, HUVECs were pre-incubated with CaD (100 µM) for 1 h. Subsequently, the cells were rinsed, VEGF_165_ (25 ng/mL) was added for 2 min. Pre-incubation of cells with CaD did not significantly reduce VEGFR-2 activation after VEGF stimulation (Fig. 1C, lane 3) suggesting that the inhibitory effect of CaD on VEGFR-2 phosphorylation is not mediated by the direct binding of CaD to cell surface components of endothelial cells.

In the third setup, both cells and VEGF_165_ were pre-incubated with CaD to investigate for possible dual effect of CaD on the ligand and the receptors. We observed an additive effect of CaD inhibition of VEGFR-2 signaling (Fig. 1C, lane 4). The inhibitory effect of CaD seems to be mainly mediated by the direct binding of CaD to VEGF and to a lesser extent interacting with the cell surface components of the endothelial membrane.

### CaD attenuates VEGF-induced phosphorylation of MEK/ERK1/2 MAP Kinase

We furthermore investigated the effect of CaD on the VEGF-induced signaling cascade (28). Treatment with VEGF_165_ induced a strong phosphorylation of the ERK1/2 MAP kinase (Fig. 2 lane 5). Co-treatment with CaD (100 and 200µM) attenuated VEGF_165_-induced phosphorylation of ERK1/2 by 40% (Fig. 2, lanes 7-8). The MAPK kinase MEK1/2 is known to be the direct upstream kinase of the ERK1/2 MAP kinase. We next investigated the effects of CaD on this signaling molecule upstream of the ERK1/2 MAP kinase, CaD markedly attenuated VEGF_165_ stimulated phosphorylation of MEK1/2 (Fig. 2).

**Fig. 2.**
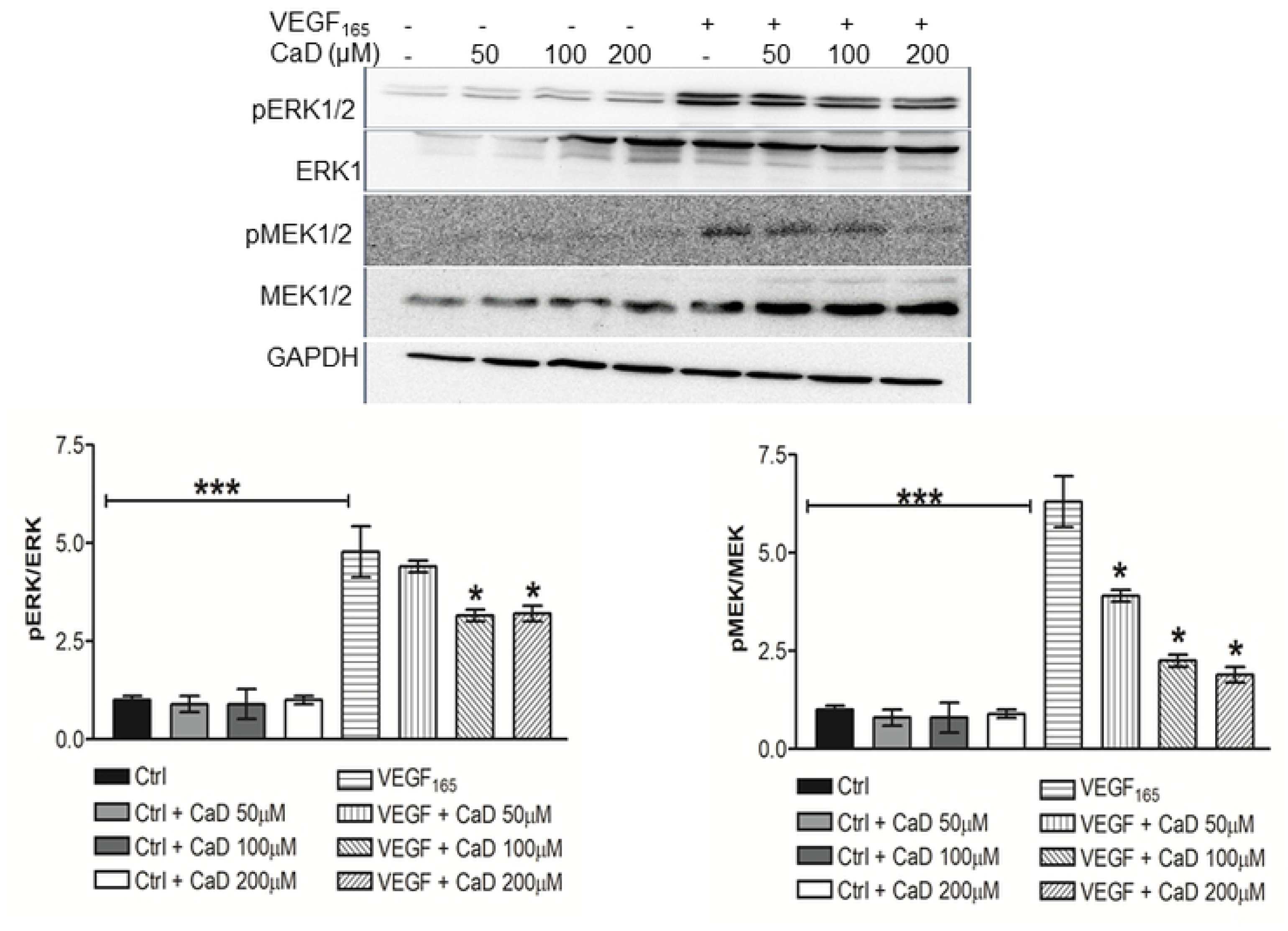
Calcium dobesilate (CaD) inhibits VEGF-induced MEK/ERK MAP kinase activation. CaD at 50, 100 and 200µM was incubated with VEGF (25 ng/mL) for 60 min before exposure to HUVECs for 15 min. Phosphorylated ERK and MEK1/2 was determined by Western blot as described in Figure 2. Data are expressed as mean ± SD (n = 3). * p < 0.05, *** p < 0.001, significantly different from VEGF-treated HUVECs.

### CaD inhibits VEGF-induced angiogenic activity in HUVECs

Treatment with CaD significantly inhibited VEGF-induced proliferation and migration of HUVECs (Fig. 3A-C). Peripheral accumulation of F-actin was detected only in VEGF-stimulated cells (Fig. 3D middle panels). Treatment with CaD completely abrogated VEGF induced accumulation of peripheral actin-rich lamellipodia-like structures (lower panels).

**Fig. 3.**
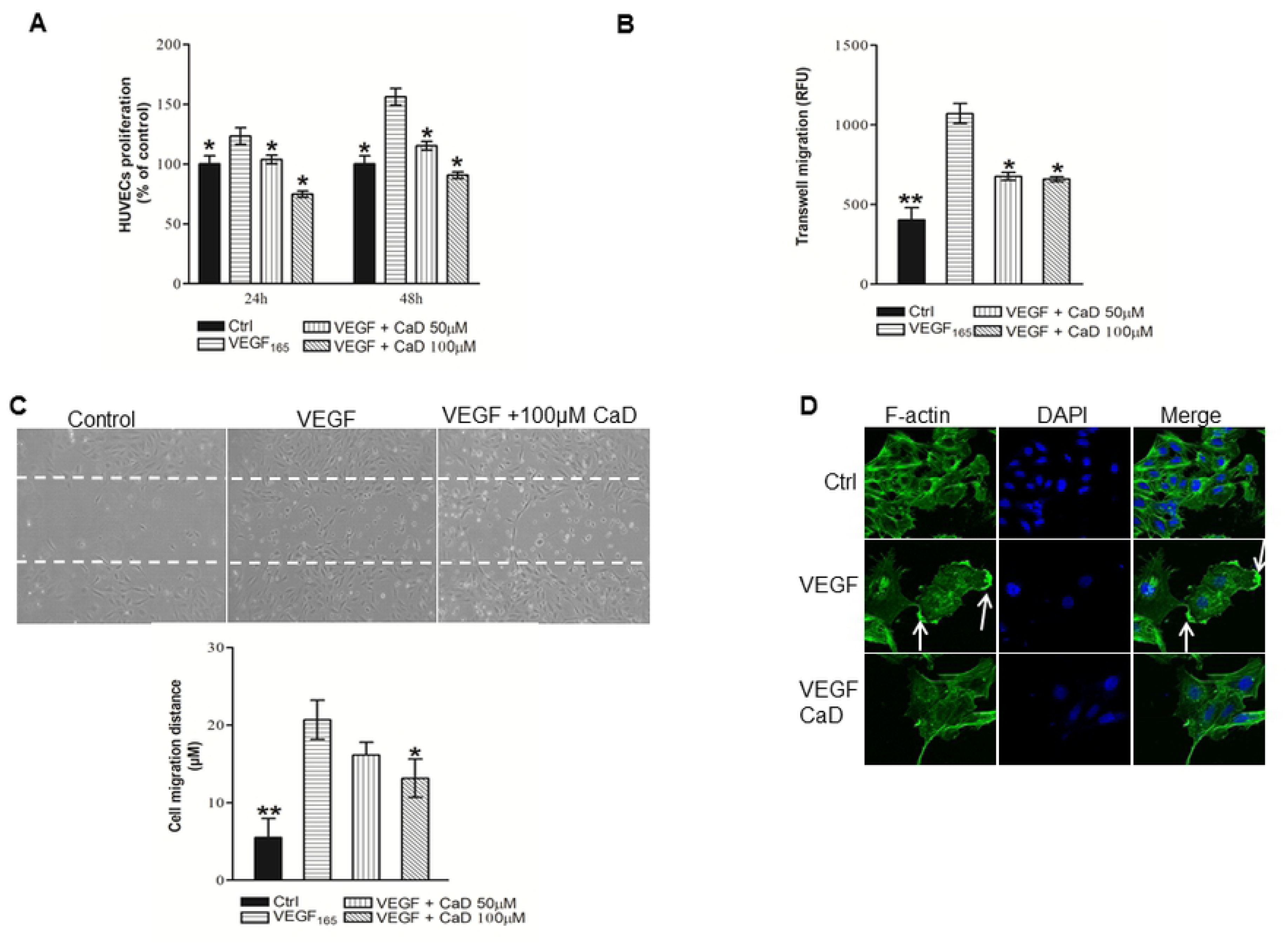
Inhibition of endothelial cell proliferation, invasion and migration by CaD. HUVECs were plated in 96-well plates, allowed to attach overnight, and then cultured for 24 h-48 h with CaD in the presence of VEGF_165_ (20ng/ml)-CaD mixture (A) at the indicated concentration. The proliferation was measured as described in the Materials and Methods section. (B) Serum-starved HUVECs were allowed to migrate through trans-well membranes towards a vehicle, VEGF_165_ (20ng/ml) and/or CaD at the indicated concentration for 24 h. Cells that had migrated to the underside of the membrane were processed for calcein-AM staining as described in the materials and methods section. (C) A monolayer of HUVECs was scratched and fresh medium containing vehicle, VEGF and/or CaD was then added. After 14 h, migration distance of HUVECs was quantified. Original magnification, ×40. (D) HUVECs were incubated with vehicle, VEGF_165_ (20ng/ml) and CaD for 10 min. Cells were stained with phalloidin-Alexa Fluor488 (left images) and DAPI (middle images). Merged view is represented in the right images. Original magnification, ×40. The results shown are the means ± SD of four independent experiments conducted in triplicate. *P<0.05, **P<0.01, versus VEGF_165_-treated HUVECs.

### Effect of CaD on VEGF_165_-induced tight junction disruption and permeability

The tight junction proteins occludin, claudin-5 and ZO-1 were expressed by HUVECs as shown by Western blot analysis. Treatment with CaD significantly prevented the decrease in ZO-1, Occludin and claudin-5 expression induced by VEGF (Fig. 4A, lanes 5 and 6). The expression of ZO-1 and claudin-5 proteins was dependent on the concentration of CaD. To test for the effect of CaD on endothelial permeability, VEGF/CaD was added to the basolateral sides of HUVECs confluent monolayers, cells were cultured for an additional 2 hours, and then processed for dextran permeability. VEGF treatment caused a significant increase in FITC-dextran flux which was significantly decreased by CaD treatment (Fig. 4B).

**Fig. 4.**
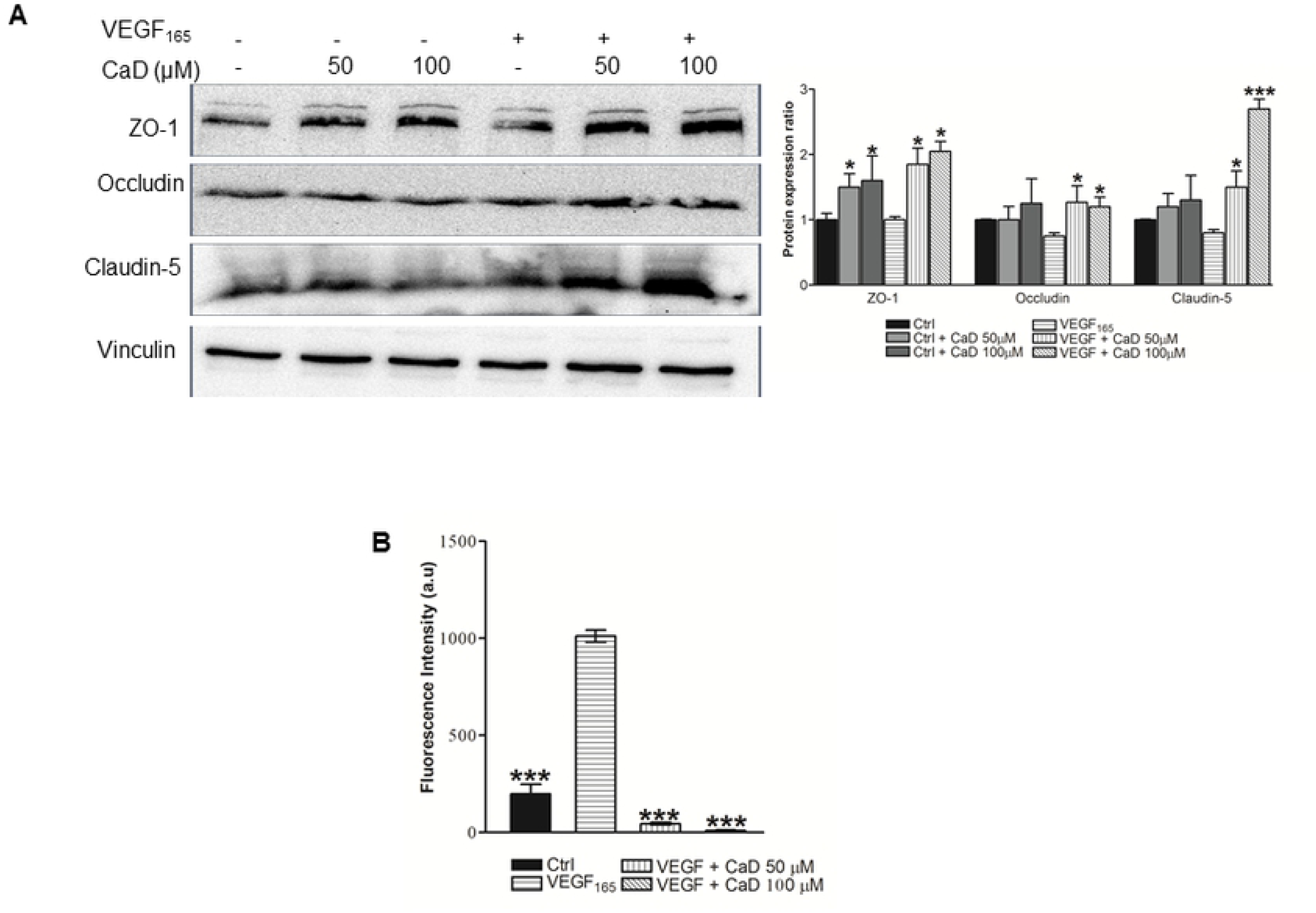
CaD prevents the decrease in tight junction protein levels and the increase in HUVECs permeability induced by VEGF_165_. Tight junction protein levels were determined by Western blotting (A) and endothelial cells permeability was by FITC-Dextran assay (B) as described in material and methods section. A representative Western blot is shown and each bar indicates the relative expression to that of VEGF_165_ alone. The average of three independent experiments ± SD. *P<0.05, ***P<0.001.

### Mechanism of action of CaD inhibitory effect on VEGF_165_

CaD (100 µM), VEGF (25 ng/ml) and heparin (10 µg/ml) were premixed for 1 h before addition to the HUVECs for 2 min. Phosphorylation of VEGFR-2 was examined by Western blot analysis. As revealed by Western blot analysis, the VEGF_165_-induced phosphorylation of VEGFR-2 increased in the presence of heparin (Fig. 5A, lane 5). Heparin abrogated CaD inhibitory effect on VEGFR-2 phosphorylation (Fig. 5A, lane 6). For comparison, we examined the effects of CaD on VEGF_121_, an isoform without heparin binding domain (HBD) (29). CaD did not significantly inhibit VEGF_121_-induced tyrosine phosphorylation of VEGFR-2 (10%) (Fig. 5B, lanes 5 and 6).

**Fig. 5.**
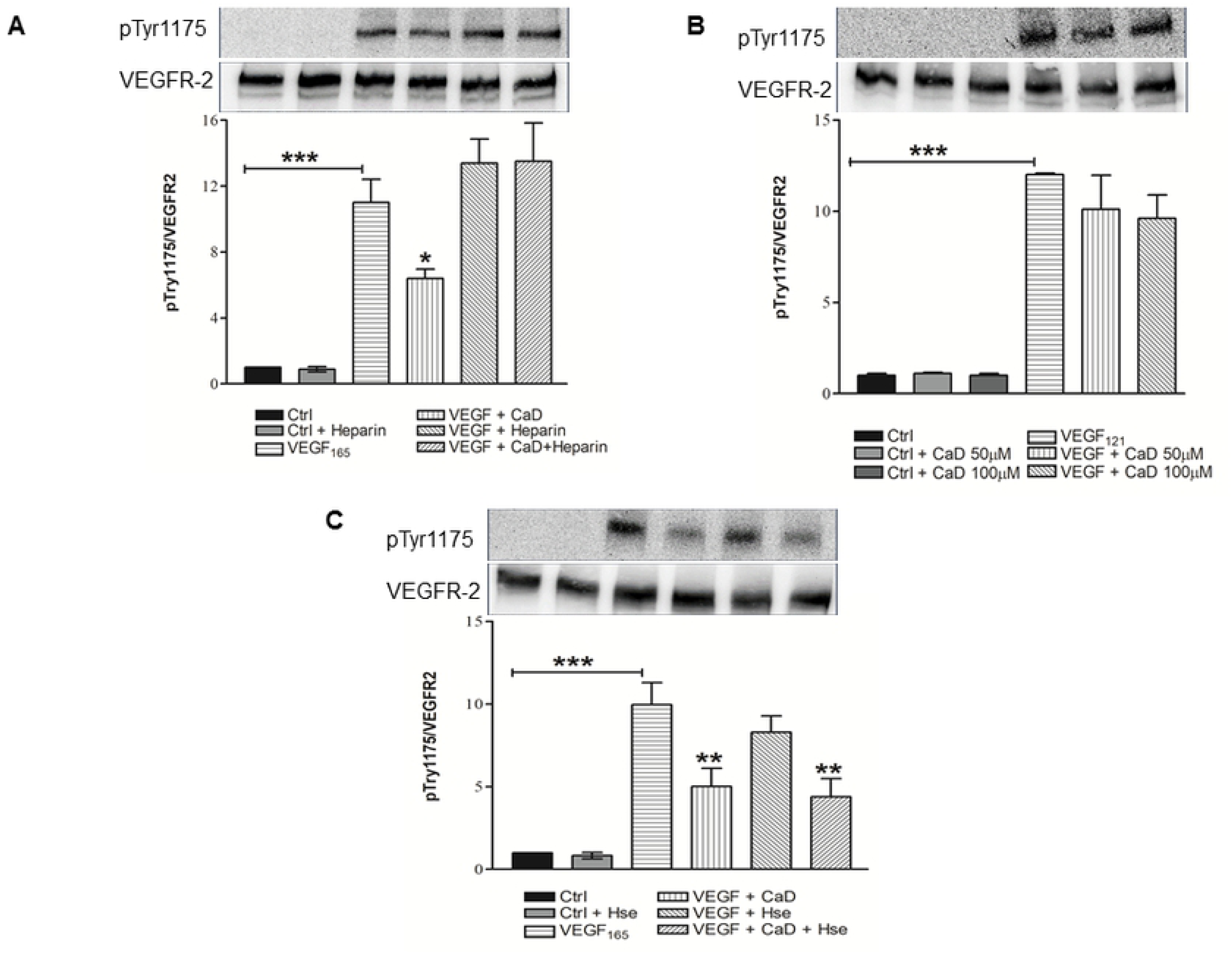
Effect of heparin and heparinase (Hse) treatment on the inhibitory effect of CaD. (A) VEGF_165_ (25 ng/mL), CaD (100µM) and heparin (10µg/ml) were pre-mixed and incubated at 37°C for 60 min prior to HUVECs stimulation for 2 min. (B) HUVECs were stimulated with VEGF_121_(25ng/ml)-CaD mixture as described in figure 1 for 2 min. (C) HUVECs were digested with heparinase (0.5µg/ml). After 30 min, each dish was washed and then stimulated with 25ng/ml VEGF_165_ alone or VEGF-CaD (100µM) mixture for 2 min. Samples were subjected to gel electrophoresis, and Western blotting was performed as described. Each bar indicates the relative phosphorylation to that of VEGF alone and the average of at least three independent experiments ± SD. *P<0.05, **P<0.01, ***P<0.001.

### CaD inhibits VEGF_165_-induced phosphorylation of VEGFR-2 in heparinase treated cells

Previous studies showed that the capacity of VEGF_165_ to bind its receptors on endothelial cells was abolished by heparinase treatment and that the effect of heparinase could be reversed by the addition of heparin (30,31). Cells were treated with or without heparinase (30 min), then washed and stimulated with CaD-VEGF_165_ for 2 min. VEGF_165_-induced phosphorylation of VEGFR-2 was reduced to 75% by heparinase treatment (Fig. 5C lane 5) suggesting that VEGF-induced receptor phosphorylation is dependent in part on the presence of the heparan sulfates (30,32). Interestingly, the decrease in VEGF_165_-induced phosphorylation by digestion with heparinase was further decreased to 50% by CaD (Fig. 5C, lane 6).

### CaD inhibits formation of VEGF_165_-VEGFR-1/2 or VEGF-heparan sulfate complexes

Because both VEGF-A_165_ and VEGFR1/2 bind heparin (33,34), exogenous heparin may also play a cross-bridging role in the engagement of the protein ligand with its receptor and CaD could destabilize this complex. We performed an *in vitro* cell-free solid phase binding assay for both VEGFR-1 and VEGFR-2. CaD concentration-dependently inhibited biotinylated-VEGF_165_ (bt-VEGF_165_) binding to VEGFR-1/2 (Fig. 6A). In conformity to the previous studies, heparin increased the binding of bt-VEGF_165_ to VEGFR-1/2 at lower concentrations (0.01-1 µg/ml) (32,35), whereas higher heparin concentrations (10-1000 µg/ml) inhibited bt-VEGF binding to the receptors (Data not shown). In the presence of heparin (1 µg/ml), CaD inhibitory effect on bt-VEGF_165_ binding to VEGFRs is abrogated (Fig. 6B). As expected Duolink *in situ* proximity ligation assay (PLA) further confirmed the inhibited interaction between VEGF-VEGFR-2 and between VEGF-HS in the presence of CaD (Fig. 6C). Quantification of the PLA signal revealed a 60% decrease in VEGF-VEGFR-2 and VEGF-HS complexes in the presence of CaD but only 12% decrease in HS-VEGFR-2 complexes (Fig. 6D). Our data further suggest that CaD interferes with HS binding to the ligand and not to the receptors.

**Fig. 6.**
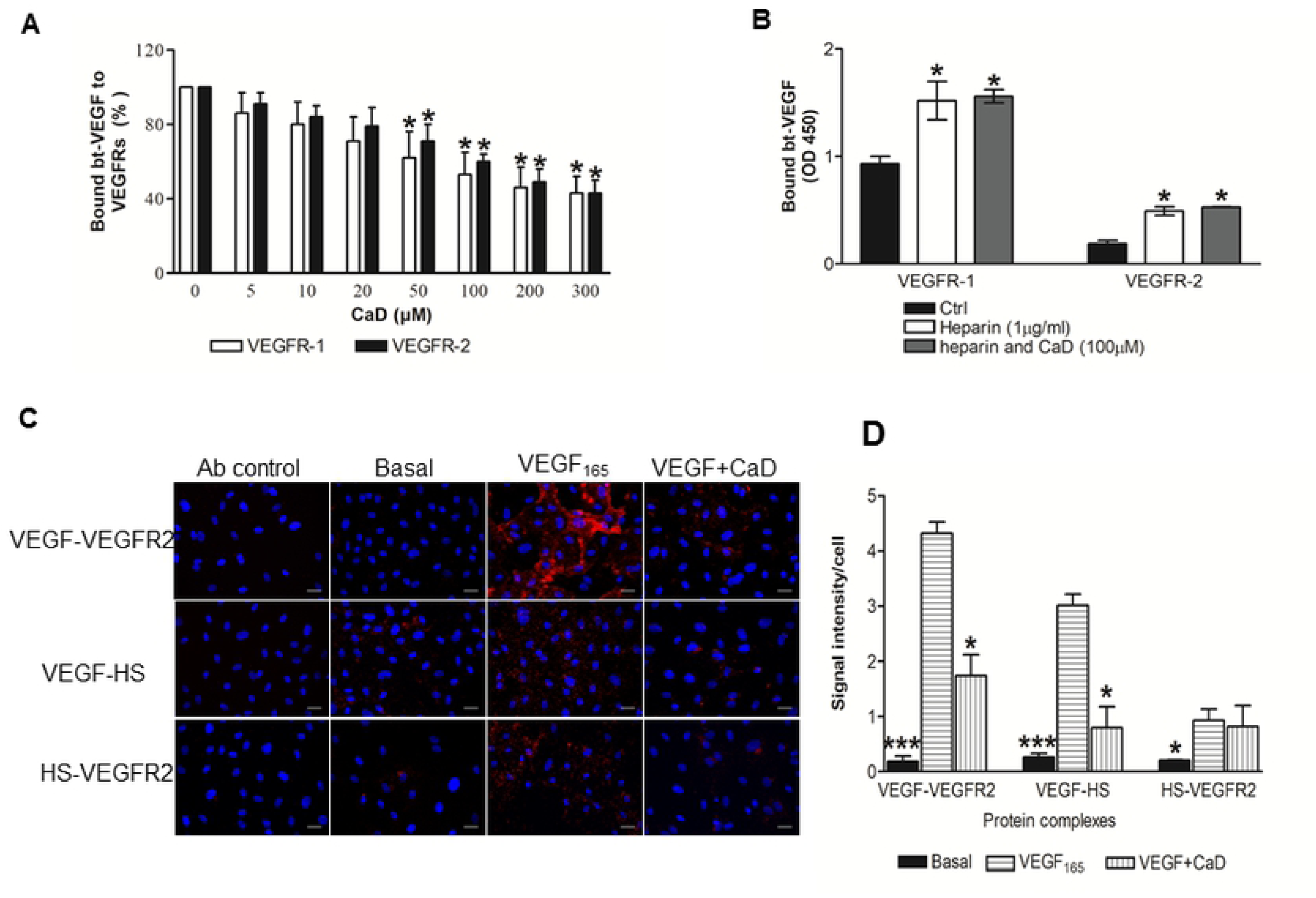
Effect of CaD on VEGF165 binding to VEGFR-1/2, heparin and HS. (A) bt-VEGF_165_ binding assay to immobilized VEGR-1/2 in the presence of increasing concentrations of CaD. Data are expressed as percentage of control (no CaD) ± SD in binding from four independent experiments; *p < 0.05 vs. no CaD. (B) bt-VEGF165 binding in the absence or presence of 1 μg/ml heparin or CaD (100 µM) to VEGFR-1 and VEGFR-2. Data are expressed as mean OD at 450 nm ± SD from four independent experiments; *p < 0.05 vs. no heparin or CaD (Ctrl). (C) Representative images were obtained by proximity ligation assay. Cells were fixed and incubated with antibodies to human VEGF (goat IgG) and to human VEGFR-2 (rabbit IgG) (upper panel) or to VEGF and heparin sulphate (mAb 10E4) (middle panel) or to VEGFR-2 and HS (lower panel) followed by proximity ligation assay reagents. Each red dot indicates a protein interaction. Nuclei are shown in blue. The antibody (Ab) control panel represents cells incubated with the corresponding IgG. (D) Quantified data presented as red dots (signal intensity)/cell. The error bar represents SD. n = 3 separate experiments *P<0.05, ***P<0.001 relative to that of VEGF alone. All images were taken with a Leica DMI3000 B microscopy, scale bar 100 µM and analyzed with NIH ImageJ software.

### Protective effects of CaD treatment in a type I diabetes mouse model

The effect of CaD compared to enalapril treatment was further investigated in vivo using type I diabetes mouse model (STZ-induced diabetes). Treatment with CaD/enalapril had no effect on glucose levels and body weight in diabetic mice (Fig. 7A-C). CaD but not enalapril significantly reduced diabetic nephropathy as reflected by serum creatinine levels (Fig. 7D) and albuminuria (Fig. 7E). CaD treatment also reduced diabetic neuropathy. After 6 week diabetes a reduction of the sensory nerve conduction velocity was observed in STZ/vehicle and STZ/enalapril groups but not in STZ/CaD group. At week 12 decreased nerve conduction velocity was observed in all STZ groups compared to nondiabetic controls, but in the STZ/CaD group the decrease was significantly less pronounced (Fig. 7F).

**Fig. 7.**
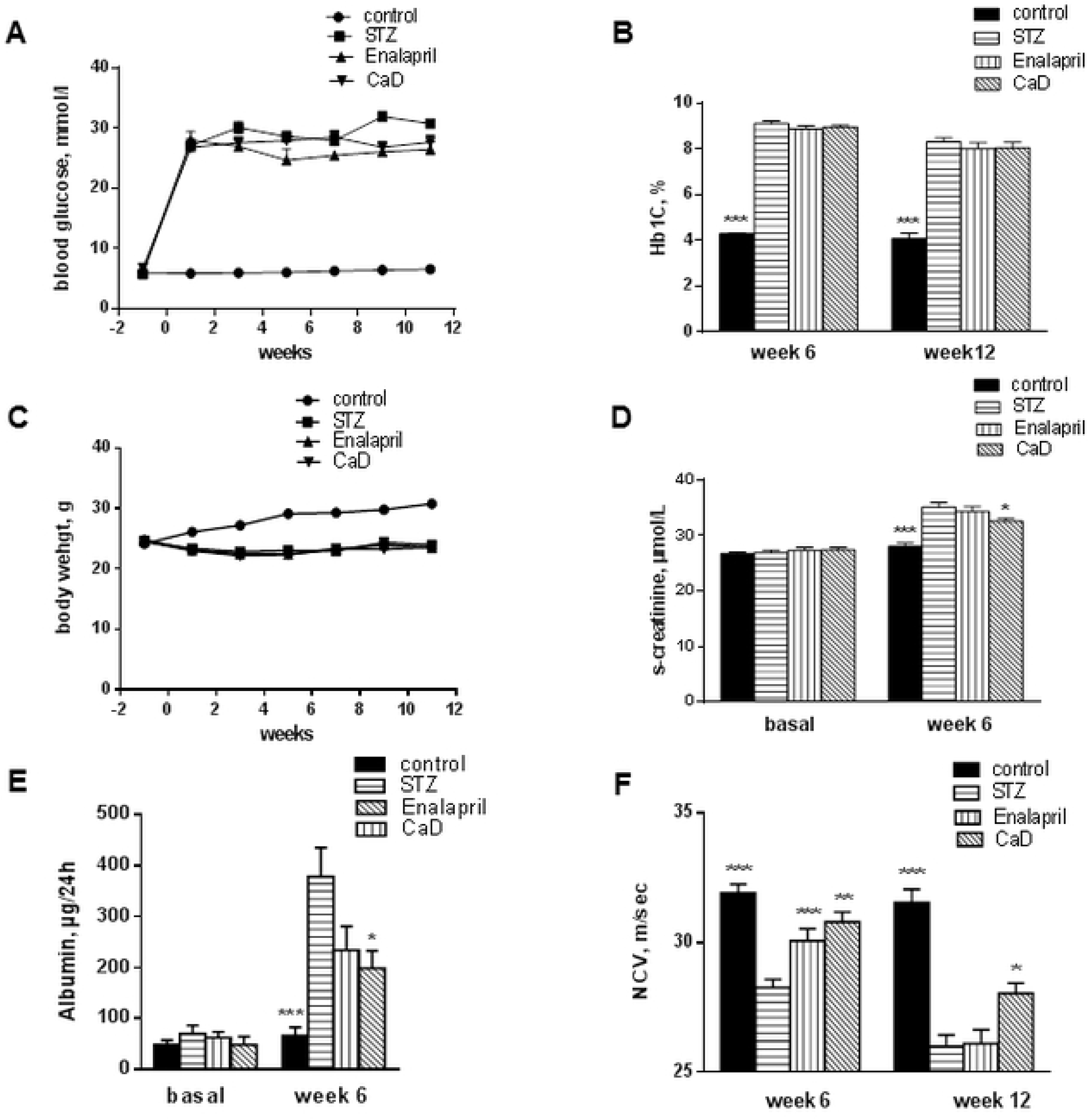
Effect of CaD treatment on diabetic complications in vivo. Type I diabetes was induced in Sv129 mice and thereafter the diabetic mice were daily treated with vehicle or CaD (100mg/kg) or enalapril (30mg/kg). CaD treatment had no effect on hyperglycemia (A and B) and body weight (C). CaD treatment protects kidney function in diabetic mice as reflected by serum creatinine levels (D) and albuminuria (E). CaD treatment reduces diabetic neuropathy as reflected by sensory nerve conduction velocity measurements (F). The results are expressed as the mean+SEM (n=20 mice for each group). *p<0.05, **p<0.01 and ***p<0.001 versus vehicle treated diabetic mice (STZ).

We next investigated the effects of CaD on diabetes-induced VEGF signaling in the kidney. CaD/enalapril treatment decreased p-VEGFR2 level in the kidney compared to the vehicle-treated diabetic mice (Figure 8A) and significantly suppressed diabetes-induced ERK1/2 (Fig. 8B, lane 4 and column 4) and P38 phosphorylation (Fig. 8C). It was accompanied by reduced inflammation in the diabetic kidneys as reflected by prevented up-regulation of CXCL-1, IL-1ß, TNF-α and IL-6 expression (Fig.9A). Whereas both CaD and enalapril were effective for these four cytokines, only CaD but not enalapril inhibited significantly MCP-1 up-regulation (Fig.9A). Moreover, diabetes-induced up-regulation of VEGF in the kidney was down-regulated by CaD/enalapril treatment (Fig. 9B). In line with reduced levels of pro-inflammatory mediators, an increased amount of F4/80 positive macrophages in the interstitial areas of diabetic kidney was significantly reduced in CaD treated mice (Fig. 9C). Additionally, CaD treatment prevented the diabetes-induced mesangial proliferation and glomerulosclerosis (Fig. 9D).

**Fig. 8.**
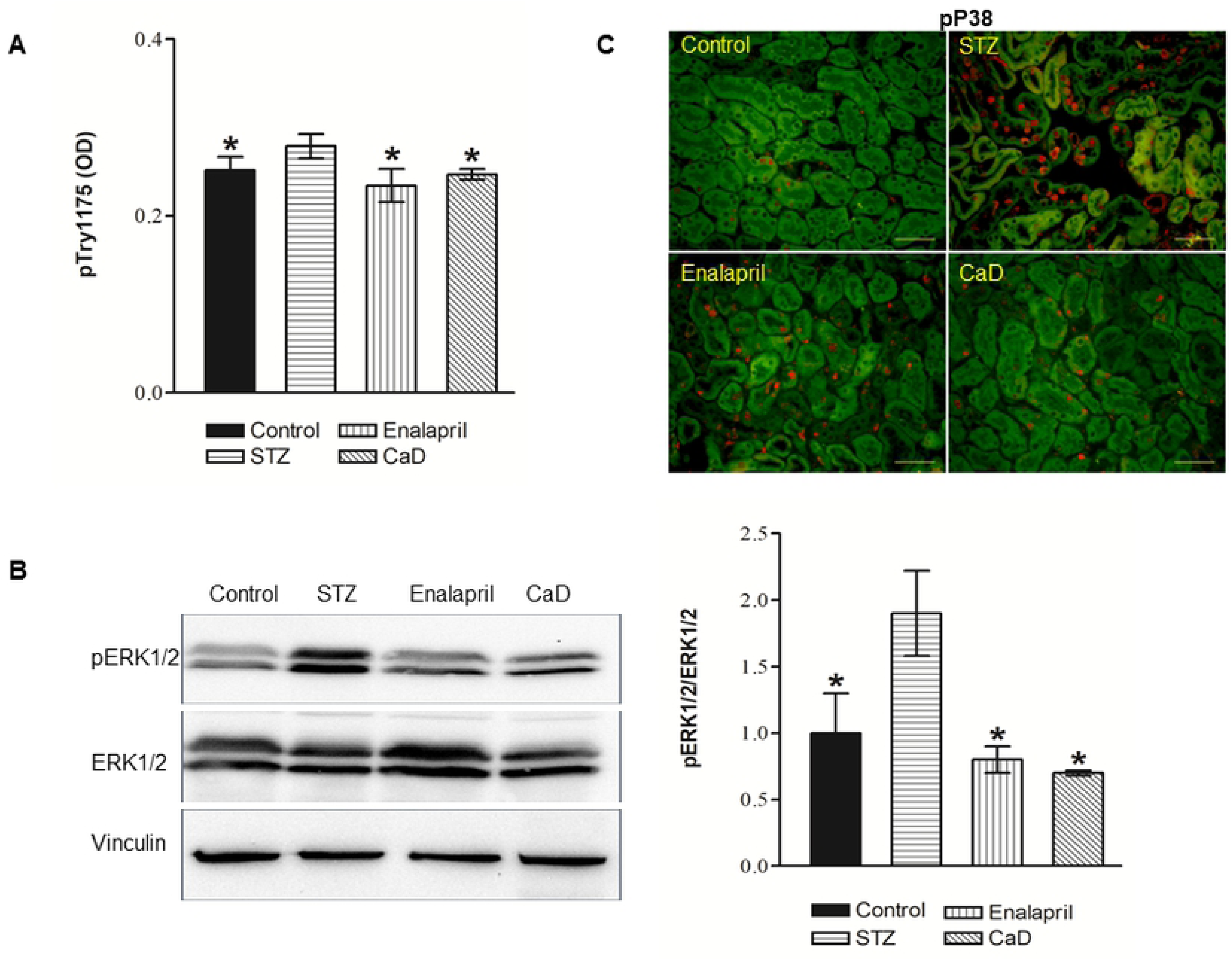
CaD inhibits VEGFR-2 phosphorylation and signaling in STZ diabetic mice. STZ mice were daily treated with vehicle, CaD or Enalapril and scarified at day week 12 for the analysis. Non diabetic mice were used as control. Kidneys were isolated and homogenized as described in materials and methods. Phosphorylated VEGFR-2 (A) in the kidney lysates was determined by ELISA mean ± SD of 5-6 animals. Phosphorylated ERK1/2 (B) was analyzed by Western blot as described in figure 1. Vinculin was used as a loading control. Phosphorylated P38 (C) was analyzed by immunohistochemistry as described in the materials and methods section. Autofluorescence is shown in green, scale bar 100 µM. Representative image for n=6/condition is depicted. *P<0.05 versus vehicle treated STZ mice

**Fig. 9.**
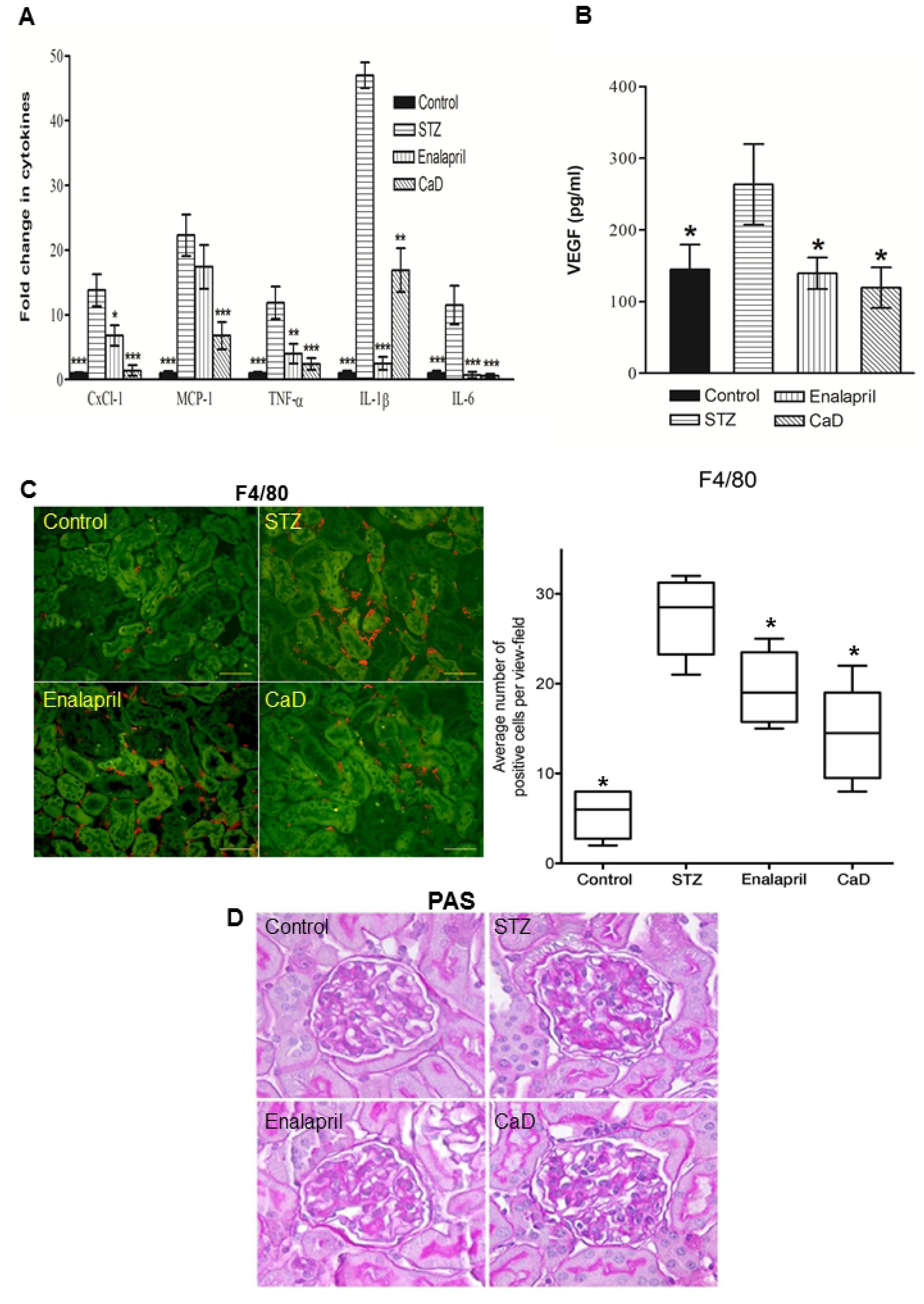
CaD prevents the increase in diabetes-induced renal inflammation and glomerulosclerosis. The mRNA levels were assessed by real time PCR and normalized to Actin (A). VEGF production was determined by VEGF_164-188_ ELISA according to the manufacturer’s instructions (B). Data represent the mean ± SD of 4-8 animals. * p < 0.05. Significantly different from vehicle treated STZ mice. Immunohistochemistry for F4/80 (red) in kidney cross sections of non-diabetic (control), diabetic vehicle treated (STZ), enalapril and CaD treated STZ mice (C). Autofluorescence is shown in green, scale bar 100 µM. Images are exemplary for n = 5/condition. Infiltrating cells were quantitatively analyzed by scoring five areas in each kidney section for F4/80-positive cells. Each bar represents the mean ± SD.* p < 0.05, **P<0.01, *** p < 0.001. Significantly different from vehicle treated STZ mice. (D) Representative kidney sections (magnification x400) stained by periodic acid Schiff (PAS) after 12 weeks of treatment (n=5).

## Discussion

CaD is used in Asia and South America to treat diabetic retinopathy (36), chronic venous insufficiency, and various conditions associated with excessive angiogenesis (21,22). Recent studies suggested that CaD exerts protective effects against diabetic nephropathy (37). Despite its broad use, the pharmacology of CaD has received little attention. We showed that CaD significantly blocked VEGF and diabetes-induced VEGFR-2 phosphorylation, which is the main mediator of proliferation, migration, survival, and permeability in endothelial cells (38). Earlier studies showed that the MAPK signaling cascade, ERK and P38, were also modulated via VEGFR-2 signaling activation by VEGF on HUVECs (39). We therefore, investigated the effect of CaD on VEGF and diabetic-induced ERK1/2 and P38 phosphorylation in HUVECs and in our mouse model respectively. We also monitored the effects on VEGF-induced endothelial cell proliferation, invasion, and migration and found these components relevant for novel blood vessel formation to be significantly inhibited in the presence of CaD. Furthermore, we found that CaD significantly down-regulated VEGF_165_-induced phosphorylation of ERK1/2 and diabetic-induced phosphorylation of ERK1/2 and P38. Similar results have been reported, in which CaD significantly inhibited FGF-induced ERK phosphorylation in glioma cells and P38 in diabetic retinopathy (40,41).

Consistent with the anti-migratory function, CaD also abolished VEGF-induced polymerization of actin in lamellipodia-like structures. Zhou and colleagues have recently demonstrated an inhibition of endothelial cell proliferation and migration by CaD under hyperglycemic conditions (42), which they attributed to the corresponding changes in VEGF expression. Instead, we demonstrate a direct effect of CaD on VEGF signaling. Our findings complement those of Angulo et al., who demonstrated a significant reduction in VEGF-induced HUVECs proliferation by CaD (28). These findings support an important role for dobesilate in vascular angiogenesis.

Dysfunction of the endothelial tight junction is crucial for the development of endothelial hyper-permeability (43). We tested the effect of VEGF and CaD on endothelial cells tight junction proteins and permeability. CaD significantly restored VEGF-induced suppression of ZO-1, Occludin and Claudin-5 expression and VEGF-induced increased permeability, indicating that the protective effects could be related to stabilizing tight junction proteins.

We propose that the dominant mechanism of CaD resulting in the inhibition of VEGF-induced VEGFR-2 activation and signaling is related to the interaction between CaD and HS. Earlier studies demonstrated that CaD interferes with heparin binding on FGF and inhibits the signaling of FGF via its receptors FGFRs (23). We treated HUVECs with a mixture of VEGF_165_, CaD, and heparin and found that the inhibitory effect of CaD on VEGFR-2 phosphorylation was abrogated by the addition of heparin. By using an *in vitro* cell-free and proximity ligation assays, we show that CaD clearly interfered with the binding of VEGF to its receptors and also to HS, these findings suggest that, binding of CaD to VEGF probably lowers the affinity of VEGF to its cognate receptors because of change in three-dimensional structure of VEGF at its receptor recognition site, and/or dissociating the receptor-growth factor signaling complex as previously suggested for FGF (23). The effect of CaD was accordingly overridden by the addition of heparin. To further substantiate the involvement of heparin binding domain in the interaction with CaD, HUVECs were stimulated with VEGF_121_, an isoform without the exon-7-encoded region, which has no capacity to bind to heparin. As expected, CaD did not significantly inhibit VEGF_121_ induced receptor phosphorylation. We suggest that CaD forms a complex with VEGF_165_ and VEGFR-2, thereby inhibiting VEGF_165_-dependent signaling. HS/heparin has been proposed to regulate VEGF biological activity by binding VEGF directly (44), and also by interacting with receptors (33,34). A report by Fernandez and colleagues demonstrated the dual inhibitory action of CaD in endothelial cells by binding to both FGF and its receptors (23). Our results suggest that the possible mechanism of the CaD action is related to interaction with heparin binding VEGF_165_ and to a lesser extent to the VEGF receptors. This interpretation is further supported by our findings using non-heparin binding VEGF_121_, where CaD did not significantly affect VEGF_121_-induced VEGFR-2 phosphorylation and also by our PLA where CaD did not prevent HS-VEGFR-2 complex formation.

To further substantiate our findings, the role of cell surface HS in VEGF_165_ activity was assessed by the reduction of VEGF_165_-induced phosphorylation of VEGFR-2 in the cells digested with heparinase (30). Interestingly, CaD treatment further reduced VEGF-induced phosphorylation of the receptor in heparinase treated cells to 50%, similar to the level observed in CaD treatment without heparinase. In agreement with our above findings, these results suggest that in addition to CaD inhibiting VEGF binding to surface HS, there is a percentage of inhibition which is contributed by CaD inhibiting VEGF binding to VEGFR-2 that has to be taken into account.

We propose a CaD-induced mechanism of action involving VEGF_165_ inhibition (Fig. 10). Cell surface HS regulates VEGF_165_ binding to VEGFR-2 and VEGF_165_-dependent phosphorylation of VEGFR-2 via binding to the heparin-like domain. In the presence of CaD, VEGF_165_, VEGFR-2 complex formation with cells surface heparan-sulfate proteoglycans is abrogated resulting into an unstable complex which is either fast degraded or VEGF_165_ binds to its receptors with low affinity. As a result, VEGF_165_-induced signaling processes, such as the phosphorylation of VEGFR-2, are decreased by CaD. It is also possible that in addition to binding to the ligand, CaD binds albeit with low significance to the HBD of VEGFR-2 partially blocking VEGF_165_ from interacting with its receptors and therefore, contributing to signaling inhibition.

**Fig. 10.**
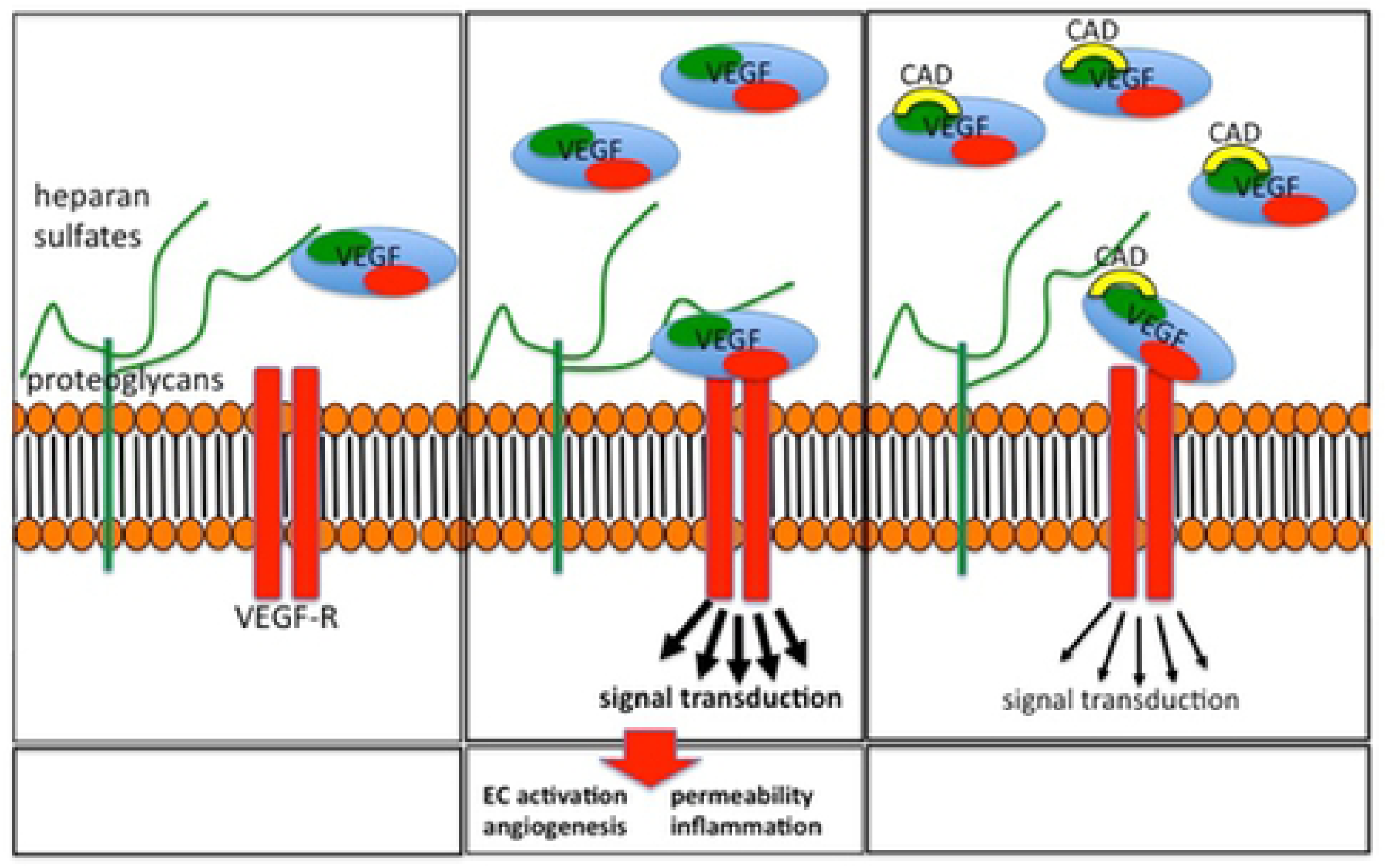
Postulated model of interactions between VEGF_165_, VEGFR-2, and CaD. VEGF_165_ binds to its co-receptor heparin sulfates (HS) of the endothelial glycocalyx with a specific binding site (left box) which stabilizes the VEGF-VEGF-R binding leading to phosphorylation of VEGFR-2 receptor, intracellular signaling and cell activation (middle box). CaD interacts with the heparin-binding domain of the VEGF_165_ (right box), thereby displacing HS from its binding site, and decreases VEGF-induced intracellular signaling. CaD also regulates VEGF_165_ activity by participating in the formation of unstable VEGF-VEGFR-2 complex.

Our proposed mechanism explains not only the CaD inhibitory effect on VEGF, but also the low rate of adverse effects as compared to VEGF antibodies in diabetes (23,45). While VEGF antibodies completely block the effects of VEGF on the intracellular signaling pathways and thereby also block VEGF-induced signals which are necessary for endothelial cell survival, interference with the heparan sulfate binding sites reduces the binding of VEGF to its receptor and, therefore, reduces its effects on endothelial cells but does not abolish the effect of VEGF on its specific membrane-bound receptor.

The severity of nephropathy is usually defined by proteinuria, which is closely correlated to renal damage. Following treatment with CaD, renal function was improved significantly as evidenced by decreased serum creatinine and urinary albumin. In addition, our experiments show that CaD prevents the increase and upregulation/activation of several pro-inflammatory cytokines (IL-6, CXCL1, IL-1ß, TNF-α, and MCP-1) that play a significant role in renal-disease progression. Although the anti-inflammatory properties of CaD have been previously reported (42,46), to the best of our knowledge, our report is the first involving a diabetic-nephropathy animal model. These findings are therefore important due to the pivotal role of inflammation in the pathogenesis of diabetic nephropathy.

CaD inhibited VEGF-induced endothelial permeability and could protect against blood-brain-barrier leakage. Such an effect was observed in the retina correlated with a decrease in the levels of VEGF in the retina (47), however, was explained by a direct effect of CaD on VEGF expression (42). Although we also observed down-regulation of VEGF production in the kidneys from STZ mice, we believe that this is a secondary phenomenon after improvement of endothelial cell function.

Interestingly, we observed a strong effect of CaD on diabetic neuropathy. Although not in similar settings, these results could be extrapolated to those of Sola-Adell et al., (47). They reported retinal neuroprotective effect of CaD in a diabetic mouse model. Further studies are required to demonstrate the effectiveness of CaD in the treatment of peripheral diabetic neuropathy. Since CaD pharmacokinetics is already know (48), and it is currently used to treat vascular complications of diabetic retinopathy (49), our findings demonstrated the therapeutic potential of CaD in the early stages of DN given that at present, anti-VEGF antibody or tyrosine kinase inhibitors therapy for diabetic nephropathy is not warranted (50).

In summary, we demonstrated that CaD inhibits VEGF signaling and function in endothelial cells and that this effect is mediated via a novel mechanism interfering with the complex formation between VEGF, VEGF-R and HS. We could also show that CaD ameliorates diabetic nephropathy in a streptozotocin-induced diabetic mouse model by VEGF inhibition. We suggest a novel mechanism to interfere with VEGF signaling and suggest that the class of CaD compounds should be investigated further, particularly in the pathogenesis of diabetic nephropathy.

## Acknowledgements

We thank Frank Hausadel, Petra Wübbolt-Lehmann and Birgit Habermeir for excellent technical assistance.

## References

1. Flyvbjerg A, Dagnaes-Hansen F, De Vriese AS, Schrijvers BF, Tilton RG, Rasch R. Amelioration of long-term renal changes in obese type 2 diabetic mice by a neutralizing vascular endothelial growth factor antibody. Diabetes. 2002;51(10):3090–4.

2. Group TDC and CT of DI and CR. Retinopathy and Nephropathy in Patients with Type 1 Diabetes Four Years after a Trial of Intensive Therapy. N Engl J Med. 2000;342(6):381–9.

3. Parving HH, Andersen AR, Smidt UM, Hommel E, Mathiesen ER, Svendsen PA. Effect of antihypertensive treatment on kidney function in diabetic nephropathy. Br Med J (Clin Res Ed). 1987;294(6585):1443–7.

4. Bakris GL, Ritz E. The message for World Kidney Day 2009: Hypertension and kidney disease - A marriage that should be prevented. Vol. 27, Journal of Hypertension. 2009. p. 666–9.

5. Wu HY, Huang JW, Lin HJ, Liao W-CC, Peng Y-SS, Hung KY, et al. Comparative effectiveness of renin-angiotensin system blockers and other antihypertensive drugs in patients with diabetes: systematic review and bayesian network meta-analysis. BMJ. 2013;347:f6008.

6. Collins AJ, Foley RN, Chavers B, Gilbertson D, Herzog C, Johansen K, et al. US renal data system 2011 Annual data report. Am J Kidney Dis. 2012;59(1 SUPPL. 1):6386.

7. Nelson RG, Newman JM, Knowler WC, Sievers ML, Kunzelman CL, Pettitt DJ, et al. Incidence of end-stage renal disease in Type 2 (non-insulin-dependent) diabetes mellitus in pima indians. Diabetologia. 1988;31(10):730–6.

8. Calles-Escandon J, Cipolla M. Diabetes and endothelial dysfunction: A clinical perspective. Vol. 22, Endocrine Reviews. 2001. p. 36–52.

9. Cooper ME, Vranes D, Youssef S, Stacker SA, Cox AJ, Rizkalla B, et al. Increased renal expression of vascular endothelial growth factor (VEGF) and its receptor VEGFR-2 in experimental diabetes. Diabetes. 1999;48(11):2229–39.

10. Braun L, Kardon T, Reisz-Porszasz ZS, Banhegyi G, Mandl J. The regulation of the induction of vascular endothelial growth factor at the onset of diabetes in spontaneously diabetic rats. Life Sci. 2001;69(21):2533–42.

11. Hohenstein B, Hausknecht B, Boehmer K, Riess R, Brekken RA, Hugo CPM. Local VEGF activity but not VEGF expression is tightly regulated during diabetic nephropathy in man. Kidney Int. 2006;69(9):1654–61.

12. Wang W, Merrill MJ, Borchardt RT. Vascular endothelial growth factor affects permeability of brain microvessel endothelial cells in vitro. Am J Physiol. 1996;271(6 Pt 1):C1973–80.

13. De Vriese AS, Tilton RG, Elger M, Stephan CC, Kriz W, Lameire NH. Antibodies against vascular endothelial growth factor improve early renal dysfunction in experimental diabetes. JAmSocNephrol. 2001;12(1046–6673):993–1000.

14. Ku DD, Zaleski JK, Liu S, Brock TA. Vascular endothelial growth factor induces EDRF-dependent relaxation in coronary arteries. Am J Physiol. 1993;265(2 Pt 2):H586–92.

15. Nakagawa T, Kosugi T, Haneda M, Rivard CJ, Long DA. Abnormal angiogenesis in diabetic nephropathy. Diabetes. 2009;58(7):1471–8.

16. Barleon B, Sozzani S, Zhou D, Weich HA, Mantovani A, Marmé D. Migration of human monocytes in response to vascular endothelial growth factor (VEGF) is mediated via the VEGF receptor flt-1. Blood. 1996;87(8):3336–43.

17. Schrijvers BF, Flyvbjerg A, Tilton RG, Lameire NH, De Vriese AS. A neutralizing VEGF antibody prevents glomerular hypertrophy in a model of obese type 2 diabetes, the Zucker diabetic fatty rat. Nephrol Dial Transplant. 2006;21(2):324–9.

18. Lai X-X, Xu R-A, Yu-Ping L, Yang H. Risk of adverse events with bevacizumab addition to therapy in advanced non-small-cell lung cancer: a meta-analysis of randomized controlled trials. Onco Targets Ther. 2016;9:2421–8.

19. Eremina V, Quaggin SE. Biology of anti-angiogenic therapy-induced thrombotic microangiopathy. Semin Nephrol. 2010;30(6):582–90.

20. Eremina V, Jefferson JA, Kowalewska J, Hochster H, Haas M, Weisstuch J, et al. VEGF Inhibition and Renal Thrombotic Microangiopathy. N Engl J Med. 2008;358(11):1129–36.

21. Cuevas P, Arrazola JM. Therapeutic response of rosacea to dobesilate. Eur J Med Res. 2005;10(10):454–6.

22. Cuevas P, Arrazola JM. Dobesilate in the treatment of plaque psoriasis. Eur J Med Res. 2005;10(9):373–6.

23. Fernández IS, Cuevas P, Angulo J, López-Navajas P, Canales-Mayordomo Á, González-Corrochano R, et al. Gentisic acid, a compound associated with plant defense and a metabolite of aspirin, heads a new class of in vivo fibroblast growth factor inhibitors. J Biol Chem. 2010;285(15):11714–29.

24. Thamm K, Njau F, Van Slyke P, Dumont DJ, Park J-K, Haller H, et al. Pharmacological Tie2 activation in kidney transplantation. World J Transplant. 2016;6(3):573.

25. Martins-Green M, Petreaca M, Yao M. Chapter 8 An Assay System for In Vitro Detection of Permeability in Human “Endothelium.” Vol. 443, Methods in Enzymology. 2008. p. 137–53.

26. Narayanaswamy PB, Tkachuk S, Haller H, Dumler I, Kiyan Y. CHK1 and RAD51 activation after DNA damage is regulated via urokinase receptor/TLR4 signaling. Cell Death Dis. 2016;7(9).

27. Goncalves V, Gautier B, Garbay C, Vidal M, Inguimbert N. Development of a chemiluminescent screening assay for detection of vascular endothelial growth factor receptor 1 ligands. Anal Biochem. 2007;366(1):108–10.

28. Angulo J, Peiró C, Romacho T, Fernández A, Cuevas B, González-Corrochano R, et al. Inhibition of vascular endothelial growth factor (VEGF)-induced endothelial proliferation, arterial relaxation, vascular permeability and angiogenesis by dobesilate. Eur J Pharmacol. 2011;667(1–3):153–9.

29. Edward Conrad H, Edward Conrad H. Chapter 2 – Structures of Heparinoids. In: Heparin-Binding Proteins. 1998. p. 7–60.

30. Ashikari-Hada S, Habuchi H, Kariya Y, Kimata K. Heparin Regulates Vascular Endothelial Growth Factor 165-dependent Mitogenic Activity, Tube Formation, and Its Receptor Phosphorylation of Human Endothelial Cells. J Biol Chem. 2005;280(36):31508–15.

31. Gitay-Goren H, Soker S, Vlodavsky I, Neufeld G. The binding of vascular endothelial growth factor to its receptors is dependent on cell surface-associated heparin-like molecules. J Biol Chem. 1992;267(9):6093–8.

32. Wijelath E, Namekata M, Murray J, Furuyashiki M, Zhang S, Coan D, et al. Multiple mechanisms for exogenous heparin modulation of vascular endothelial growth factor activity. J Cell Biochem. 2010;111(2):461–8.

33. Park M, Lee ST. The fourth immunoglobulin-like loop in the extracellular domain of FLT-1, a VEGF receptor, includes a major heparin-binding site. Biochem Biophys Res Commun. 1999;264(3):730–4.

34. Dougher AM, Wasserstrom H, Torley L, Shridaran L, Westdock P, Hileman RE, et al. Identification of a heparin binding peptide on the extracellular domain of the KDR VEGF receptor. Growth Factors. 1997;14(4):257–68.

35. Nishiguchi KM, Kataoka K, Kachi S, Komeima K, Terasaki H. Regulation of pathologic retinal angiogenesis in mice and inhibition of VEGF-VEGFR2 binding by soluble heparan sulfate. PLoS One. 2010;5(10).

36. Zhang XY, Liu W, Wu SS, Jin JL, Li WH, Wang NL. Calcium dobesilate for diabetic retinopathy: a systematic review and meta-analysis. Sci China Life Sci. 2014;58(1):101–7.

37. Yang W, Yu X, Zhang Q, Lu Q, Wang J, Cui W, et al. Attenuation of streptozotocin-induced diabetic retinopathy with low molecular weight fucoidan via inhibition of vascular endothelial growth factor. Exp Eye Res. 2013;115:96–105.

38. Holmes K, Roberts OL, Thomas AM, Cross MJ. Vascular endothelial growth factor receptor-2: Structure, function, intracellular signalling and therapeutic inhibition. Vol. 19, Cellular Signalling. 2007. p. 2003–12.

39. Shibuya M. Vascular Endothelial Growth Factor (VEGF) and Its Receptor (VEGFR) Signaling in Angiogenesis: A Crucial Target for Anti-and Pro-Angiogenic Therapies. Genes and Cancer. 2011;2(12):1097–105.

40. Cuevas P, Díaz-Gonzalez D, García-Martín-Córdova C, Sánchez I, Lozano RM, Giménez-Gallego G, et al. Dobesilate diminishes activation of the mitogen - activated protein kinase ERK1/2 in glioma cells. J Cell Mol Med. 2006;10(1):225–30.

41. Leal EC, Martins J, Voabil P, Liberal J, Chiavaroli C, Bauer J, et al. Calcium dobesilate inhibits the alterations in tight junction proteins and leukocyte adhesion to retinal endothelial cells induced by diabetes. Diabetes. 2010;59(10):2637–45.

42. Zhou Y, Yuan J, Qi C, Shao X, Mou S, Ni Z. Calcium dobesilate may alleviate diabetes-induced endothelial dysfunction and inflammation. Mol Med Rep. 2017;16(6):8635–42.

43. Liu E, Morimoto M, Kitajima S, Koike T, Yu Y, Shiiki H, et al. Increased Expression of Vascular Endothelial Growth Factor in Kidney Leads to Progressive Impairment of Glomerular Functions. J Am Soc Nephrol. 2007;18(7):2094–104.

44. Leung D, Cachianes G, Kuang W, Goeddel D, Ferrara N. Vascular endothelial growth factor is a secreted angiogenic mitogen. Science (80-). 1989;246(4935):1306–9.

45. Loges S, Mazzone M, Hohensinner P, Carmeliet P. Silencing or Fueling Metastasis with VEGF Inhibitors: Antiangiogenesis Revisited. Vol. 15, Cancer Cell. 2009. p. 167–70.

46. Bogdanov P, Solà-Adell C, Hernández C, García-Ramírez M, Sampedro J, Simó-Servat O, et al. Calcium dobesilate prevents the oxidative stress and inflammation induced by diabetes in the retina of db/db mice. J Diabetes Complications. 2017;31(10):1481–90.

47. Solà-Adell C, Bogdanov P, Hernández C, Sampedro J, Valeri M, Garcia-Ramirez M, et al. Calcium Dobesilate Prevents Neurodegeneration and Vascular Leakage in Experimental Diabetes. Curr Eye Res. 2017;42(9):1273–86.

48. Tejerina T, Ruiz E. Calcium dobesilate: Pharmacology and future approaches. Vol. 31, General Pharmacology. 1998. p. 357–60.

49. Garay RP, Hannaert P, Chiavaroli C. Calcium dobesilate in the treatment of diabetic retinopathy. Vol. 4, Treatments in Endocrinology. 2005. p. 221–32.

50. Tanabe K, Maeshima Y, Sato Y, Wada J. Antiangiogenic Therapy for Diabetic Nephropathy. Biomed Res Int. 2017;2017:1–12.

